# Cultivation of the polyextremophile *Cyanidioschyzon merolae* 10D during summer conditions on the coast of the Red Sea and its adaptation to hypersaline sea water

**DOI:** 10.1101/2023.02.02.526792

**Authors:** Melany V. Villegas, Ricardo E. González-Portela, Bárbara Bastos de Freitas, Abdulaziz Al Jahdali, Gabriel I. Romero-Villegas, Raghdah Malibari, Rahul Vijay Kapoore, Claudio Fuentes-Grünewald, Kyle J. Lauersen

## Abstract

The west coast of Saudi Arabia borders the Red Sea, which maintains high average temperatures and increased salinity compared to other seas or oceans. Summer conditions in the Arabian Peninsula may exceed the temperature tolerance of most currently cultivated microalgae. The Cyanidiales are polyextremophilic red algae whose native habitats are at the edges of acidic hot springs. *Cyanidioschyzon merolae* 10D has recently emerged as an interesting model organism capable of high-cell density cultivation on pure CO_2_ with optimal growth at 42 °C and low pH between 0.5-2. *C. merolae* biomass has an interesting macromolecular composition, is protein rich, and contains valuable bio-products like heat-stable phycocyanin, carotenoids, β-glucan, and starch. Here, photobioreactors were used to model *C. merolae* 10D growth performance in simulated environmental conditions of the mid-Red Sea coast across four seasons, it was then grown at various scales outdoors in Thuwal, Saudi Arabia during the Summer of 2022. We show that *C. merolae* 10D is amenable to cultivation with industrial-grade nutrient and CO_2_ inputs outdoors in this location and that its biomass is relatively constant in biochemical composition across culture conditions. We also show the adaptation of *C. merolae* 10D to high salinity levels of those found in Red Sea waters and conducted further modeled cultivations in nutrient enriched local sea water. It was determined that salt-water adapted *C. merolae* 10D could be cultivated with reduced nutrient inputs in local conditions. The results presented here indicate this may be a promising alternative species for algal bioprocesses in outdoor conditions in extreme desert summer environments.

## 1 Introduction

Currently, bioprocesses that use microalgae are being implemented with increasing intensity in nearly all countries due to their broad potential for low input waste-revalorization. Algae can convert the nutrients and trace elements from waste streams into their biomass using light energy via photosynthesis and CO_2_ as a carbon source (Fuentes-Grünewald et al., 2021). There are numerous market opportunities for different products made from algal biomass such as animal feed, food, pigments, bioplastics, speciality chemicals, and biostimulant fertilizers (Rumin et al., 2020). The worldwide production of microalgae biomass is currently low, ~56,000 tons per annum compared to production of marine seaweeds, ~34.6M tons per annum (Ferdouse et al., 2018; Cai et al., 2021). There is currently an opportunity, especially in desert countries, for increased algal production processes aimed at bio-resource circularity, especially connected to urban and industrial areas.

Algae can be cultivated in photobioreactors built on non-arable land and directly connected to industrial side-streams to derive value from effluents produced in otherwise overlooked geographical areas such as desert regions or other extreme environments. The implementation of algal bioprocesses in growing contexts will further promote algal biomass and bioproduct integration into common use. Microalgal bioprocesses are most strategically suited to geographies with high solar irradiation and stable temperatures, having local CO_2_ emissions sources, non-arable land, access to sea water, and inexpensive energy sources. It was recently reported that the geographical areas with the highest potential for onshore marine microalgae biomass production were between 30°N and 30°S in the so-called “hot belt countries” (Greene et al., 2022). This area is dominated by countries with enormous desert regions such as southern US (Arizona), Northern Chile (Atacama), Northern Africa, Northern Australia, and the Middle East. It is predicted that in these environments, microalgal biomass production from local inputs can be maximized. To capture the value of resources in waste-streams, the development and integration of algal bioprocesses can be parts of circular reuse concepts in these environments. New target microalgal species with high biotechnological potential, that can tolerate extreme conditions must be identified, characterized, and developed for these contexts.

The unicellular red microalgae *Cyanidioschyzon merolae* is a poly-extremophile that thrives in low pH and high temperatures. Its biomass contains interesting amounts of valuable metabolites such as thermostable phycocyanin (PC) in addition to starch, β-glucan, β-carotene and zeaxanthin carotenoid pigments. This extremophile exhibits its best growth at temperatures from 42-50 °C and low pH (0.5-2.5) (Miyagishima and Tanaka, 2021). Tolerance to acidic conditions can allow cultivation with continuous injection rates of high concentration CO_2_ and ammonia while also minimizing contamination. This species may be a promising candidate for algal bioprocesses in extreme environments. It is an excellent candidate for waste circularization concepts which seek to take high concentration feed streams, especially in summertime conditions in desert environments, and convert them into valuable biomass.

Here, we show the performance characteristics of *C. merolae* 10D in lab-scale photobioreactors modelling different seasons on the mid Red Sea coast in Saudi Arabia, successful scaling of these cultures to outdoor pilot scale (1 m^3^) cultures in summer desert conditions, biomass characterization of the strain grown outdoors, and the adaptation of this species to the high saline conditions to allow cultivation with medium made using Red Sea water. This is the first report that *C. merolae* 10D has been grown at a pilot scale to identify its potential use for industrial applications in desert contexts. Our results indicate that *C. merolae* 10D, and maybe other Cyanidiophyceae, will be valuable for bioproduction or waste-stream bio-conversion in summer conditions in extreme desert environments where other common commercial algal species may reach temperature threshold limitations.

## 2 Materials & Methods

### 2.1 Lab-scale cultivation of *C. merolae* 10D and bioreactor growth tests

*C. merolae* 10D (Toda et al., 1995; Matsuzaki et al., 2004) was received from the lab of Prof. Peter Lammers (Arizona Center for Algae Technology Innovation (AzCATI), Arizona State University (ASU)) which we routinely maintained in MA2 liquid medium (Minoda et al., 2004): pH 2.3, adjusted with H_2_SO_4_ in 125 mL Erlenmeyer flasks shaken at 100 rpm under 90 μmol photon m^−2^ s^−1^ (hereafter, *μE)* continuous white light and 42 °C in a Percival algae incubator supplemented with 4% CO_2_ in air mixtures. Preculturing prior to growth analysis was performed by inoculation of cells to shake flasks with MA2 medium for 4 days, then cells were resuspended according to desired densities in 400 mL flasks prior to growth analysis in Algem photobioreactors (Algenuity©, UK). Cultures in biological duplicates with starting cell concentrations of 3×10^6^ cells mL^−1^ were cultivated in environmental simulations of the mid Red Sea coast conditions of Thuwal, Saudi Arabia (22.3046N, 39.1022E) as recently reported (de Freitas et al., 2023) and compared with control cultures using 12:12 h light:dark or 24 h constant 1500 *μ*E illumination at 42 °C. All cultures received continuous 7% CO_2_ in air at a 25 mL min^−1^ flow rate. Modeled environmental conditions were from each season with local historical weather data used to build profiles of temperature and light for February (Winter), May (Spring), August (Summer), and November (Autumn). Daily, 15 mL of algae culture from each replicate flask was taken for cell concentration (100 μL), and biomass quantification (3x 4 mL). Cell densities were measured using an Invitrogen Attune NxT flow cytometer (Thermo Fisher Scientific, UK) equipped with a Cytkick microtiter plate autosampler unit. Each *C. merolae* 10D biological replicate was diluted 1:100 with 0.9% NaCl solution and loaded into a 96-well microtiter plate in technical duplicates prior to injection into the flow cytometer using the autosampler. Samples were mixed three times immediately before analysis, and the first 25 μL of the sample was discarded to ensure a stable cell flow rate during measurement. For data acquisition, 50 μL from each well was analyzed. Optical densities were measured as absorbance at 740 nm (OD _740nm_) by the Algem photobioreactors every 10 min. Cell dry mass was measured by vacuum filtration of 3x 4 mL of each biological replicate on pre-weighted filters (0.45μm). The algal cell masses were dried at 60 °C for 24h prior to weighing.

To adapt *C. merolae* 10D to saline conditions and enable its cultivation on a medium made from acidified seawater, MA2 medium was prepared with different dilutions of filtered and autoclaved Red Sea water (RSW-MA2) as 100, 80, 60, 40 and 20% mixtures with fresh water plus MA2 nutrients. In a 6-well plate, 200 μL *C. merolae* 10D late logarithmic phase preculture culture was inoculated into wells each containing the different RSW-MA2 concentrations. The cells were cultivated in the algal incubator for 7 days as above. Cultures which reached reasonable cell densities were re-inoculated in new media across the salinity dilution range and grown for another 7 days. This process was repeated another two times and the culture adapted to 100% RSW-MA2 was chosen for further cultivation in emulated local weather conditions in Algem photobioreactors as described above.

### 2.2 Cultivation of *C. merolae* outdoors at increasing culture volumes

Precultures of *C. merolae* 10D grown in Algem photobioreactors were pooled and transported to the Development of Algal Biotechnology Kingdom of Saudi Arabia (DAB-KSA) pilot plant (KAUST, Thuwal, Saudi Arabia), and cultivated outdoors in various photobioreactors from June-August 2022. Cultures were started in 8 L south-facing glass columns at outdoor conditions with 39 ± 4.0°C and mid-day irradiance 1503 ± 241 *μ*E, for two weeks. The culture medium was prepared using Altakamul industrial/agricultural-grade salts rather than analytical grade as in lab cultivations. The medium contained 5.3 g (NH_4_)_2_SO_4_, 1.09 g KH_2_PO_4_ and 0.481 g MgSO_4_ L^−1^ with F/2 medium trace elements added to 1 mL L^−1^ (Guillard, 1975). The pH was adjusted to 2.5 with H_2_SO_4_. Additionally, an ammonium-based booster solution (10 mL L^−1^, with 50% of the original initial concentration of NH_4_^+^) was added every two days during the whole cultivation period. Once enough inoculum was produced, the culture was transferred to a 60 L column photobioreactor (Varicon Aqua® Phyco-Bubble model) with constant CO_2_ sparging at a rate of 0.2 L min^−1^ and the unit was exposed to outdoor conditions of 32.4 ± 2 °C, 1351 ± 110 *μ*E. This culture was used as an inoculum for a 600 L raceway (Varicon Aqua® Phyco-Pond model) for 1 week, and then it was transferred to a 1,000 L tubular photobioreactor (PBR, Varicon Aqua® Phyco-Flow model) and the cultivation run for 24 days. Finally, this culture was used as inoculum for two other 1,000 L PBRs started at 30% dilution from the first reactor. Two reactors were run with 100% CO_2_ sparging - PBR-1: “Commercial” pure CO_2_ Alpha Gaz®, 99.995% (CG) and PBR-2: “Green” CO_2_ from flue gas, Gulf Cryo® (GC), while a third was not given CO_2_ - PBR-3: “Atmospheric” 0.04% CO_2_ (AC). In all treatments, gassing was supplied constantly (0.2 L min^−1^) from 8 am to 5 pm.

OD _750nm_ was recorded daily with a UV-Vis Spectrophotometer (Thermo Scientific® Genesys™ 50, UK). Cell dry weight measurement (in duplicates) was performed every other day using a gravimetric method, where aliquots were rinsed with acidified fresh water (pH 2.5) and filtered through glass microfiber filters (Whatman GF/F) and weighed after drying. Nutrient uptake was measured as N (as ammonium) and P (as phosphate), using Spectroquant® assay kits following manufacturer’s protocols. Temperature and pH were monitored continuously during the experiment (10 d) with the built-in probes of the PBRs. Light intensity was measured as irradiance (quantum flux) with a quantum meter probe (Biospherical instruments Inc® Model QSL2000). For biochemical characterizations, at the incubation site, the cell suspension was harvested and washed twice with acidified freshwater (pH 2.5). The washed microalgal pellets were freeze-dried overnight and stored in dark at −20°C until further analysis.

### 2.3 Biochemical characterization of *C. merolae* 10D biomass

All chemicals and analytical reagents were HPLC grade (Fisher Scientific, U.K.) unless stated otherwise. A multi-assay procedure (Chen and Vaidyanathan, 2012,2013; Kapoore et al., 2019) was modified for the quantification of total protein, carbohydrate, chlorophyll and carotenoids as follows. Briefly, freeze-dried microalgal pellets were weighed (1.5 - 2 mg) in 2 mL Safe-Lock microcentrifuge tubes. The dry pellets were resuspended in 24.3 μL of phosphate Buffer (pH 7.4) and 1.8 mL of 25% *(v/v)* methanol in 1M NaOH along with an equal volume of glass beads (425-600 μm i.d., acid washed). Cells were disrupted using Retsch MM 400 Mixer Mill (Retsch, GmbH) for 3 cycles (10-min bead beating and 2-min stand). For carbohydrate analysis, 2x 0.2 mL extract was transferred to 2 mL PTFE capped glass vials: for the control by adding 1.2 mL 75% H_2_SO_4_; for the experimental sample by adding 0.4 mL 75% H_2_SO_4_ and 0.8 mL anthrone reagent; completing the assay as described (Chen and Vaidyanathan, 2012, 2013; Kapoore et al., 2019; Fuentes-Grünewald et al., 2021) by incubating at 100 °C for 15 minutes followed by measurement in polystyrene cuvettes (ΔA_578_). The remainder extract after cell disruption was stored at −80 °C in 4 mL PTFE capped glass vials and later saponified as described (Chen and Vaidyanathan, 2012, 2013; Kapoore et al., 2019; Fuentes-Grünewald et al., 2021) by incubating at 100°C for 30 min. For the pigment assay, saponified extracts of 0.7 mL were transferred after vortexing to 2 mL Safe-Lock microcentrifuge tubes containing 1.05 mL 2:1 mixture of chloroform with methanol. After centrifugation and phase separation, chlorophyll and carotenoid concentrations were determined as previously described (Chen and Vaidyanathan, 2012, 2013; Kapoore et al., 2019). The remainder of the saponified extract was stored at −80 °C for subsequent protein assay where saponified extracts of 25 μL were first placed directly into 96-well assay plates with the following additions: controls, 0.2 mL BCA reagent alone (Thermo Scientific); experimental, 0.2 mL BCA/Cu mix (Thermo Scientific) and incubated at 37°C for 30 minutes, measuring ΔA_562_.

A previously reported procedure for phycocyanin extraction was modified for the assessment of this pigment in *C. merolae* 10D (Coward et al., 2016). Briefly, 1 mL of microalgal cell suspension was harvested and then centrifuged for 5 min at 2,500xg at 4 °C supernatant was decanted and the biomass was washed X3 with 1 mL of acidified water (pH 2.5). After washing, 1 mL of water was added to the wet biomass followed by four consecutive freezing (−20°C) and thawing (4°C) cycles for cell disruption. The extract was vortexed and then centrifuged at 10,000xg for 10 min to remove cell debris. The clear blue supernatant containing extracted phycocyanin was quantified using spectrophotometry with OD at 455, 564, 592, 618 and 645 nm using GENESYS 50 UV-Vis Spectrophotometer (Thermo Fisher Scientific). The concentrations of phycocyanin were quantified using the previously reported equation (Coward et al., 2016).

## 3 Results

### 3.1 Modeled growth of *C. merolae* 10D in lab-scale photobioreactors

The Cyanidiophyceae tolerate extreme conditions in their native environments on the edges of acidic hot springs. Here, we modeled whether the extreme conditions *C. merolae* 10D has evolved to thrive in would enable its cultivation in conditions which are found at the mid-Red Sea Saudi Arabian coast and by extension to similar desert countries in different seasons. Photobioreactor programs were designed using local weather-station data to set light and temperature profiles in lab-scale photobioreactors (de Freitas et al., 2023) and cultures were grown with standard laboratory MA2 medium (Minoda et al., 2004). The individual seasonal conditions, Winter - February, Spring - May, Summer - August, Autumn - November, were compared to a control culture grown with a set light cycle like the photoperiod found in Saudi Arabia (12:12 day:night), and constant temperature of 42 °C (Figure 1A). Culture performance was determined by optical (Figure 1B) and cell densities (Figure 1C) throughout the 8-day cultivation. Photographs of the cultures are also shown to indicate culture health at the end of cultivation using the different modeled seasons (Figure 1D). The best culture performance was observed in the control culture with constant temperature and OD_740nm_ above 2 were achieved fastest in Summer, Spring and Autumn reactor programs, respectively (Figure 1B).

**Figure 1.**
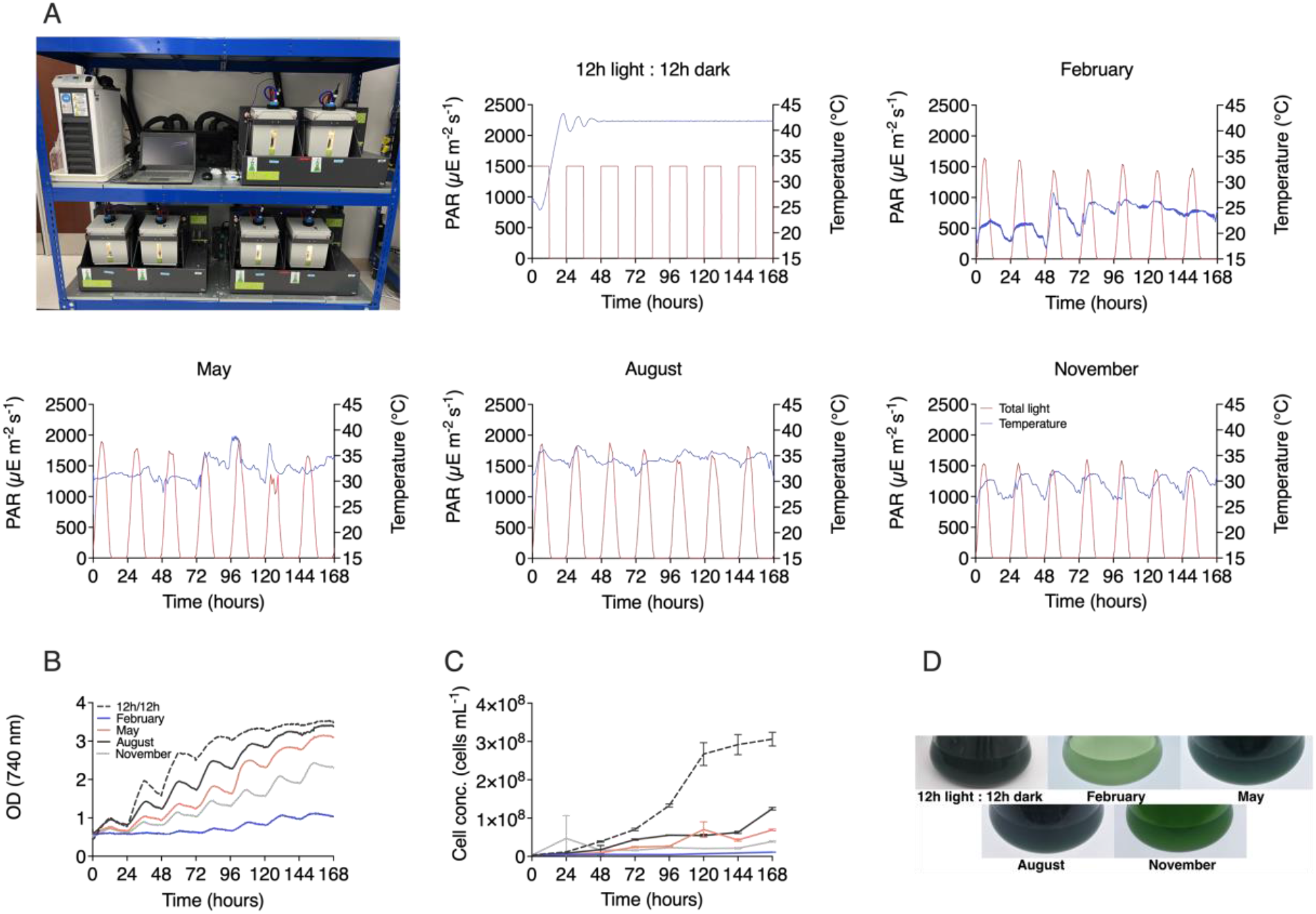
Modeled seasonal productivity variation of *C. merolae* 10D in photobioreactors. **A** temperature and light profiles in Algem photobioreactors (pictured) which are based on historical weather data from Thuwal, Saudi Arabia were used to cultivate *C. merolae* 10D in MA2 medium and assess performance across different months in this locale. A control culture was set to 42 °C with 12:12 hour day:night illumination cycling, while February, May, August, and November profiles were used to model winter, spring, summer, and autumn, respectively. Growth performance was assessed by optical density (**B**) recorded every 10 min in the bioreactors and cell densities (**C**) which were recorded daily by flow cytometry. **D** Culture health and densities were also assessed photographically at the end of cultivations.

### 3.2 Scaled cultivation of *C. merolae* 10D outdoors on the mid-Red Sea coast

From May until August 2022, scaled cultivations of *C. merolae* 10D were performed outdoors using natural light irradiance and temperature at the DAB-KSA algal pilot facility at KAUST. Cultures were taken from lab-scale cultivations above, inoculated in 8-L columns and grown in medium prepared with industrial grade fertilizers as described in the Material & Methods section. From 8-L columns (6-L working volume), *C. merolae* 10D inoculum was transferred to a 60 L vertical column with pure CO_2_ injection during daylight hours and subsequently to a 600 L raceway pond over several weeks (Figure 2). In August, the raceway pond culture was used as inoculum for starting a 1000 L tubular photobioreactor (at 0.38 g L^−1^) culture, which was maintained for 28 days of operation in batch mode (Figure 2). The culture experienced temperatures between 32-42 °C (average 37.7 °C) and daily peak irradiance of ~1800 μE (Figure 2, recorded every other day). After the 5-day lag phase, the culture exhibited a steady increase in biomass for 7 days reaching a maximum biomass production of 1.8 g L^−1^ on the 12^th^ day of cultivation (Figure 2). The maximal productivity achieved was ~300 mg L^−1^ d^−1^ between 96-288 h of the cultivation.

**Figure 2.**
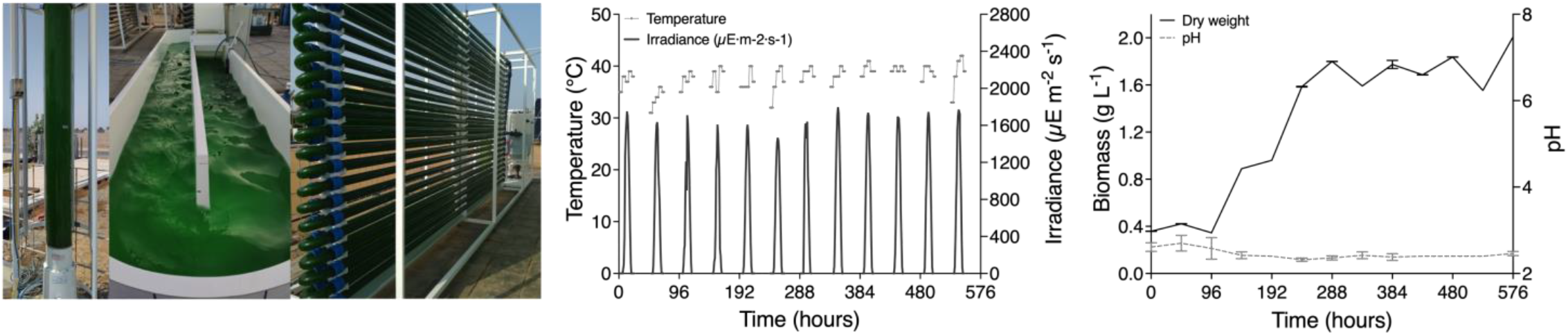
Scaled cultivation of *C. merolae* 10D in various culture set-ups in Thuwal, Saudi Arabia in July and August 2023. Cultures in a 60 L column photobioreactor were used as inoculum for a 600 L raceway pond, which was consequently used to inoculate a 1 m^3^ tubular photobioreactor (picture left). Temperature and irradiance at the tubular reactor were recorded every other day for 24 days during cultivation (middle). Culture performance was measured by cell dry weight throughout the cultivation and pH was also recorded (right).

The biomass obtained from this 1000 L tubular reactor was used as inoculum for two other reactors set beside the first (Figure 3). The inoculum allowed each reactor to be started with OD 1.0 and each reactor was cultivated for an additional 7 days (Figure 3). The reactor position determined maximal light irradiance, with the middle reactor receiving less light than the external reactors (Figure 3). Culture temperatures were relatively consistent across the three units, between 34-40 °C (average 36.5 °C) (Figure 3). One outer (east facing) and inner reactor were cultivated with pure CO_2_ injections while one outer (west facing) reactor was only sparged with air. As anticipated, the air cultivated culture (PBR 3) did not proliferate beyond inoculum density, while those given CO_2_ injections (PBR 1&2) continued to increase in their optical densities (Figure 3). These behaviors mirrored ammonium and phosphate uptake rates, which consistently was taken up by *C. merolae* cells during the cultivation period. At ~100 hours, supplemental ammonium was added to the cultures to determine if the extremophile could continue to consume this in excess, and a second boost in terms of growth (OD) was recorded. All cultures consumed the excess ammonium added (Figure 3). Regardless of CO_2_ source (beverage grade of industrially reclaimed CO_2_), the cultures given these gasses in excess were able to proliferate (Figure 3, PBR 1&2). The culture grown with air (PBR 3), exhibited higher phycocyanin content when biomass was sampled. Yet the biochemical composition of other components was consistent in samples from all reactors (Figure 3).

**Figure 3.**
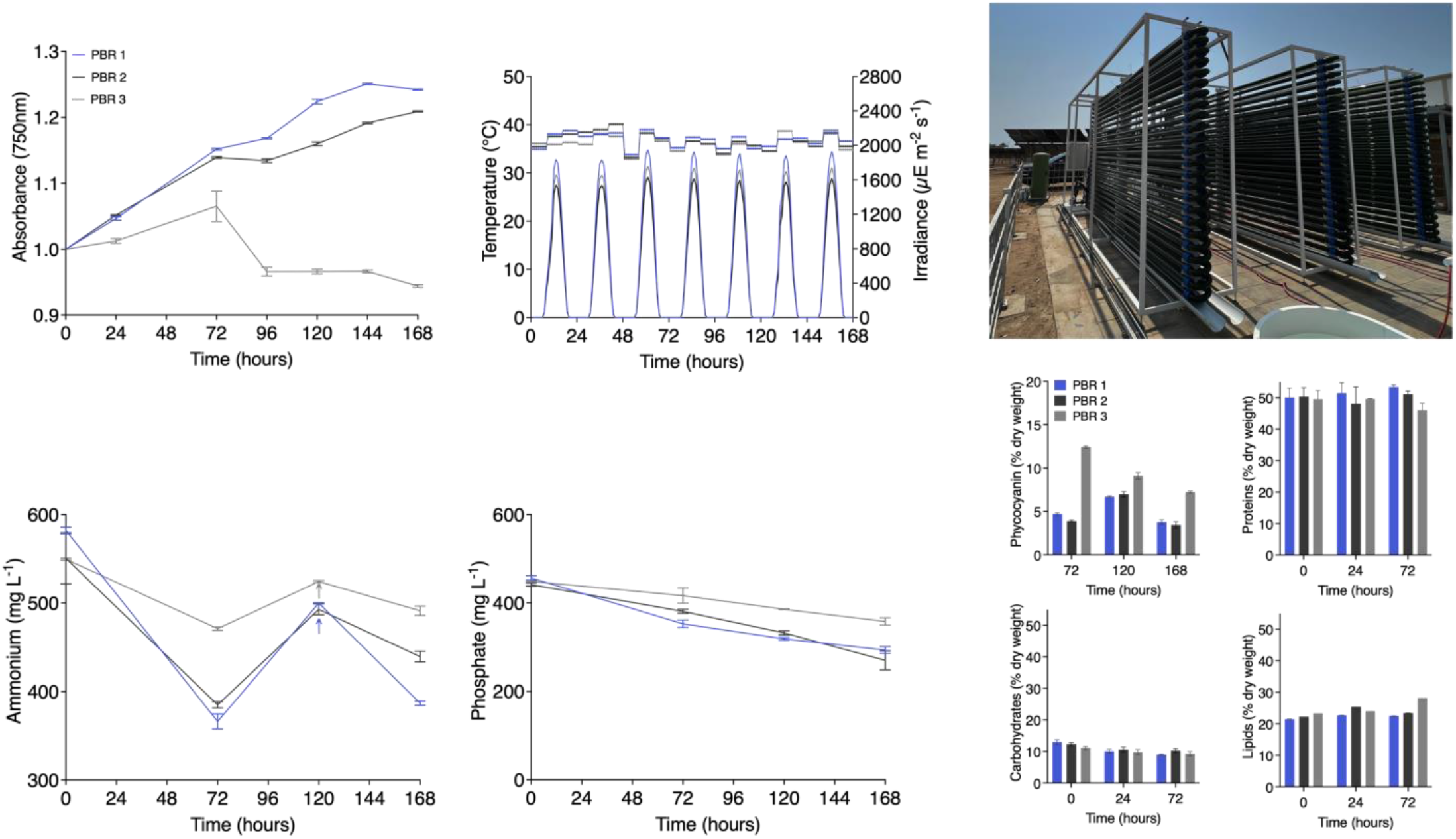
Three 1 m^3^ tubular photobioreactors were inoculated with the stationary phase culture of Figure 2. OD_750nm_, temperature, and irradiance were measured throughout the cultivation. The three reactors (pictured upper right) were given beverage grade (PBR 1), Industrially reclaimed “green” CO_2_ (PBR 2), or atmospheric air (PBR 3). The reactors each experienced slightly different irradiances based on their position to one another, with the outermost having higher incident light intensities. Ammonium and phosphate levels were recorded at different intervals, and supplemental ammonium solution was added to all reactors after 120 hours. Culture biomass was analyzed at different time-points for phycocyanin, protein, carbohydrate, and lipids content (lower right).

### 3.3 Adaptation of *C. merolae* 10D to high saline culture medium using Red Sea water

Cultivation of an extremophile in a desert environment can have the advantage of its thermotolerance, especially in summer conditions, however, the use of fresh-water resources for algal cultivation is only appropriate in these environments as part of waste-water treatment and reuse strategies. The use of seawater improves the sustainability of large scale algal cultivation concepts, especially in coastal-desert regions. It was important to determine if we could adapt the osmo-tolerance of *C. merolae* 10D to the high saline conditions found in Red Sea water as an alternative source of cultivation medium for this organism. In lab-scale photoincubators, *C. merolae* 10D was cultivated in increasing mixtures of Red Sea water supplemented with MA2 nutrients (Figure 4). Inoculum was cultivated in microtiter plates and after 7 days of light and CO_2_ conditions, cells in the best growing well were used to inoculate a fresh set of dilutions and the process repeated over several weeks (Figure 4). After 4 total passes, cells proliferating in MA2 medium made with 100 % Red Sea water were obtained (Figure 4). This adapted strain was then subjected to the same environmental modeling in photobioreactors as performed for its freshwater progenitor (Figure 1) to determine if salinity tolerance had affected its growth behavior (Figure 4). Based on optical density, the strain exhibited similar growth behaviors in saline conditions. The salt-water adapted C. *merolae* 10D performed best in constant 42 °C and 12:12 d:n control culture conditions with Summer (August) the best seasonal performance, followed by spring (Figure 4).

**Figure 4.**
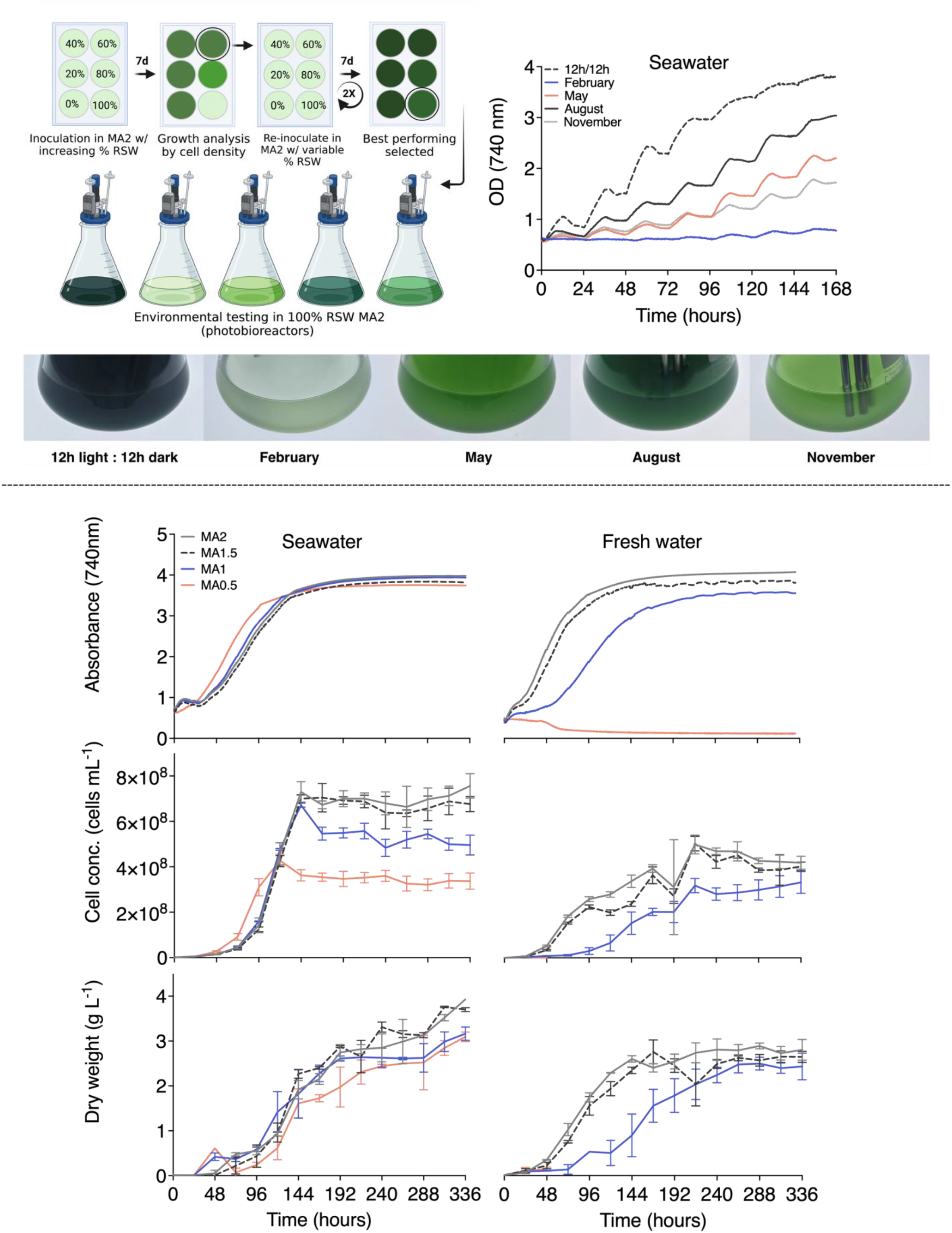
Adaptation of *C. merolae* 10D to the hypersalinity of Red Sea water. Inoculum cultures were grown for 7 days in a range of autoclaved Red Sea water mixtures with fresh water and MA2 nutrient supplementation. At the stationary phase, the salt content condition that resulted in the highest cell density was used as inoculum for the same conditions in a new culture plate. This process was repeated for a total of 4 passes until reasonable growth was observed in 100% Red Sea water MA2 medium. This culture was used as inoculum for photobioreactor modeled environmental testing of *C. merolae* 10D performance in temperature and light conditions for different seasons on the mid-Red Sea coast (as illustrated, upper left). Optical density (OD_740 nm_) was recorded continuously for the salt-water adapted *C. merolae* for a control culture (42 °C with 12:12 hour day:night illumination), modeled seasonal programs for February, May, August, and November were used to simulate winter, spring, summer, and autumn, respectively (upper right). Culture performance was also observed photographically at the end of cultivation (pictured). Saltwater adapted and its freshwater progenitor *C. merolae* 10D were subjected to continuous illumination and 42 °C temperature conditions with culture media made using different ratios of MA2 nutrient solutions to determine the possibility of reducing nutrient inputs for its growth. Culture performance was assessed by continuous optical density monitoring in the photobioreactors, daily cell density, and biomass (dry weight) measurements over 14 days (bottom panels). Figure partially created with Biorender.

As nutrients (fertilizers) can be an expensive input to algal culture concepts at large scale, and MA2 has very high concentrations of ammonium and phosphate compared to other algal cultivation media (Minoda et al., 2004), we sought to determine if it was possible to reduce the nutrient load and still achieve reasonable growth performance from *C. merolae* 10D. Here, both fresh-water and the saline-adapted *C. merolae* 10D were grown in lab-scale photobioreactors with a constant light and temperature program, but the culture medium was prepared using full, 1.5, half, and 0.5 concentrations of the nutrient solutions used in MA2 medium. Cultures were grown for 2 weeks in photobioreactors with constant light and CO_2_ gassing. Culture performance was assessed by optical and cell densities, as well as dry weights. It was determined that it is possible to dilute the nutrient composition of MA2 to MA0.5 in saline conditions and achieve a comparable performance of the alga in full medium. Red Sea water adapted *C. merolae* 10D achieved culture dry weights of 3 g L^−1^ in 14 d even when diluted to MA0.5 (Figure 4). Saline-adapted cultures grown with MA0.5 exhibited less cells per volume culture, but comparable biomass, suggesting heavier cells, this could be due to accumulation of lipids or increased starch/beta-glucan inside the cells. The optical density and biomass reached in these cultures were comparable, although slightly lower than MA1.5 and MA2 counterparts (Figure 4). However, fresh-water prepared MA2 could only be diluted to MA1 (Allen, 1959) where reduced culture performance was observed (Figure 4). Freshwater MA2 cultivated *C. merolae* 10D exhibited lower final cell densities and dry weights compared to their salt-water counterparts, although reaching comparable OD_740nm_ in a replete medium.

## 4 Discussion

### 4.1 Polyextremophilic algae for regional bioresource reuse concepts

The Arabian Peninsula is one of the more extreme environments in which human settlements have been established. Temperatures in these desert environments are consistently high, with strong irradiance and minimal precipitation (Almazroui et al., 2012; AlSarmi and Washington, 2014). These conditions are similar in most of desert countries found in the so called “hot belt” (see introduction). Despite extreme conditions, the urban population in these regions is steadily increasing, generating waste streams which require treatment and often contain nitrogen and phosphorous concentrations that can eutrophicate aquatic environments if not properly treated. This is already and issue in dense urban areas in Europe and other countries, having direct environmental impact in water bodies that cause harmful algae bloom proliferations due to nutrient emission eutrophication process. Controlled microalgal cultivation has been identified as one of the technologies to bioremediate eutrophied industrial side-streams from aquaculture, wastewater treatment plants, and anaerobic digestion facilities (Mayhead et al., 2018; Silkina et al., 2019; Fuentes-Grünewald et al., 2021). In addition, Saudi Arabia has many local sources of CO_2_ rich emissions, especially from industrial activities, that can be readily sourced with minimal transport distances. In this locale, the combination of the high irradiance, local waste-water streams, flat non-arable land and CO_2_ sources can be synergistically combined to support algal bio-processes in Saudi Arabia and neighboring countries in the Gulf Cooperation Countries (GCC) area.

The mean maximal and minimal temperatures experienced across the Arabian Peninsula are higher than the current temperate zones where microalgal cultivation is currently conducted (Figure 5). The maximal summertime temperatures exceed 45 °C, with regional variability depending on the site. These temperatures are above the threshold of heat stress and growth cessation of many currently cultivated algal species (Ras et al., 2013). In order to implement algal bioprocesses as part of broader resource circularity bio-economy drives, thermotolerant species are required for installations that will operate during the summer months in some specific countries, or throughout the year in desert countries.

**Figure 5.**
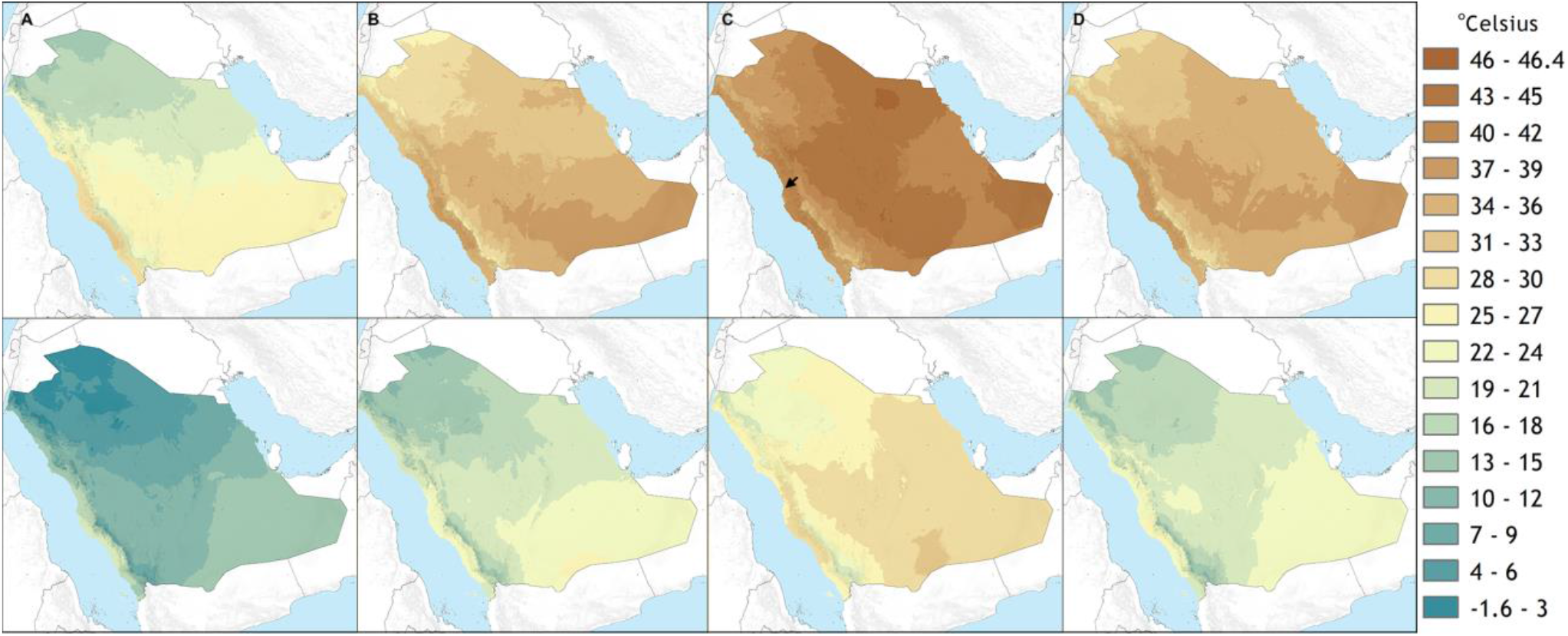
Mean maximum (upper) and minimum (lower) average monthly temperature for January (**A**), April (**B**), August (**C**), and October (**D**) at ground level from 1970-2000 in Saudi Arabia (Fick and Hijmans, 2017). Location of the KAUST DAB-KSA pilot facility site of cultivation is indicated with a black arrow.

The temperature modeling in lab-scale photobioreactors used here was designed from atmospheric weather station data at sea level, the temperatures of which would be lower than that experienced inside a photobioreactor which would warm from the solar and infrared radiation. Growth in these tests was possible in modeled August conditions but not at maximal rates of productivity observed in control cultures, and much lower when other seasons were investigated (Figure 1). *C. merolae* 10D was able to proliferate in 1000 L photobioreactors on the mid-Red Sea coast and achieve up to 2 g L^−1^ biomass at a maximal rate of 300 mg L^−1^ d^−1^ in batch mode. There, it experienced temperatures ~5 °C warmer than those modeled (Figure 2), much closer to its optimum at 42 °C. In coastal regions of this area, high humidity moderates temperature extremes compared to inland regions. It is likely that at in-land sites in urban areas, cultures would experience significantly higher temperatures than those experienced at our study site. *C. merolae* would be an ideal candidate to proliferate in such situations and is a promising strain for phased seasonal growth concepts in such an extreme environment.

### 4.2 Acidophiles in high-strength waste-stream valorization

Outdoor cultivations conducted in this study were performed with culture medium using agricultural grade chemical fertilizers. These were found to be sufficient to enable growth of the alga outdoors at various scales. Nutrients are one of the key expenses in large scale microalgae production facilities. Using agricultural grade fertilizers instead of analytical grade nutrients, the reduction in production cost, and the sustainability of microalgae production has a significant effect in the reduction of operational expenditure (Singh and Das, 2014). When the culture was grown in 3-parallel 1 m^3^ tubular reactors, the cultures tolerated addition of extra ammonium, which would cause acidic pH shifts in other algal species which require neutral pH. The cultures also tolerated either commercial beverage-grade or industrial emission reclaimed “green” CO_2_ gas sources (Figure 3). Cells grown with either CO_2_ source did not differ in biomass composition containing equivalent phycocyanin (PC), protein, carbohydrate, and lipid contents (PBR 1+2, Figure 3). The culture which was starved from carbon, only atmospheric CO_2_ levels, exhibited higher PC conentrations at 72 hours (PBR 3, Figure 3). This could be a means of increasing PC content prior to harvest, but needs further investigation. The PC of *C. merolae* is considered more thermostable than that of other currently harvest algal species and could be a valuable co-product from the biomass (Rahman et al., 2017). These behaviors suggest *C. merolae* 10D as an acidophile is highly suited to industrial waste re-valorization processes which could convert high-strength ammonia containing wastewaters and industrial CO_2_ sources into valuable biomass. We suggest this promising extremophile as a unique strain to be used in large scale production facilities for CO_2_ reuse applications.

### 4.3 Opportunities for Cyanidiophyceae biotechnology in a regional context

Extremes of heat in summer are balanced by moderate and even low temperatures in winter and spring months in the Arabian Peninsula (Figure 5). Photobioreactor modeling of C. *merolae’s* performance in these months indicated that the polyextremophile did not perform optimally under lower temperature regimes (Figure 1 and 4). From a bioprocess standpoint, engineering in-culture heating in these off-months is technically straightforward as heating processes require less energy than cooling which can be achieved, for example, with industrial heat waste (Ekendahl et al., 2018). It should be possible, therefore, to cultivate *C. merolae* 10D year-round in this environment, so long as bio-processes are designed with relevant parameters and approaches in mind.

The cultivation in acidic conditions requires some considerations in the source of inputs for scaled *C. merolae* 10D cultivations. CO_2_ injections into the culture medium (effluent inputs) could be used to reduce pH of solutions depending on alkalinity. High-strength ammonia containing waste-waters will also be appropriate for C. merolae 10D cultivation as the consumption of ammonium can will further drive the culture to acidic conditions (Henkanatte-Gedera et al., 2015; Selvaratnam et al., 2016; Nirmalakhandan et al., 2019). This additional acidification of input waters by CO_2_ may enhance the potential of *C. merolae* as a vehicle for carbon reuse, valorization, and circularity. *C. merolae* cultivation could be best performed on already acidic waste streams, like those of the dairy industry. Process designs with this organism will have to determine the best ways to incorporated its acidic cultivation conditions.

The ability to adapt *C. merolae* to saline cultivation conditions (Figure 4), also opens the possibility for cultivation in sea waters sourced along the coastlines of desert countries, improving the sustainability of commercial large scale microalgae facilities. Here, high-strength aquaculture effluents with CO_2_ injections may be used as inputs for culture of saline adapted *C. merolae* 10D. The species could be highly valuable for nitrogen and phosphorous removal from on-land marine aquaculture concepts, while generating a protein-rich biomass that can be added to feeds, generating a truly circular economy. As *C. merolae* is a cell-wall deficient species, its rupture and incorporation into feed as a protein biomass is straightforward and requires little energy inputs during downstream processing. This property also allows simple extraction of its thermostable phycocyanin or its biomass could be used to make bio-stimulant fertilizers due to its high protein content, for emerging contained environment agriculture concepts suitable for this region (Lefers et al., 2020).

## 5 Conclusions

Here, we demonstrate that the polyextremophile *C. merolae* 10D is a promising candidate for algal-based resource circularity in hot desert environments. Its thermotolerance allows its cultivation even in the extremes of desert summers and its acidic preferences can be used to minimize contamination and maximize ammonia removal from liquid and CO_2_ waste streams. The work reports scaled cultivation of C. *merolae* in the summer months in Saudi Arabia and shows that it can be adapted to salinities at least as high as those observed in Red Sea waters. C. *merolae* could be an interesting candidate for on-land marine aquaculture wastewater treatment and revalorization in addition to the reuse of high CO_2_ concentration emissions. This is the first report of the scaled cultivation of *C. merolae* 10D and the first demonstration of its growth in the Middle Eastern context. Our report sets a foundation for increasing investigations into the use of *C. merolae* and its biomass for bioresource circularity and applications like feed, fertilizer, and other high value bio-products such as thermostable phycocyanin. This report indicates that *C. merolae* 10D may hold unique promise for biotechnological application using the resources found in abundance in the Arabian Peninsula and other desert regions with urban and industrial development.

## 6 Conflict of Interest

The authors declare that the research was conducted in the absence of any commercial or financial relationships that could be construed as a potential conflict of interest.

## 7 Author Contributions

MVV was responsible for cultivation of *C. merolae* 10D in lab-scale, adaptation to sea water, lab-scale photobioreactor operation, sampling, data analysis, and figure preparation. BBdF was responsible for lab-scale bioreactor operation, sampling, data analysis, and figure preparation. REGP and GIRV were responsible for outdoor cultivation of C. merolae from 5L-1000L. RVK and RM were responsible for biochemical characterization, data interpretation, data reporting and analysis. CFG and KJL were responsible for project design, funding acquisition, data analysis and manuscript writing. All authors contributed to the writing of this manuscript and figure layout decisions.

## 8 Funding

The research reported in this publication was supported by KAUST baseline funding awarded to KL and the DAB-KSA project funded by the Saudi Ministry of Environment Water and Agriculture (MEWA) through Beacon Development (KAUST).

## 9 Acknowledgements

The authors would like to thank Prof. Peter Lammers and Dr. Mark Seger for providing Cm strain to KJL. The authors are grateful to Abhishekh Palaparambil Vijayan for GIS data visualization used in Figure 5.

## 11 Supplementary Material

Supplementary Data File 01. Data used to make Figure 1

Supplementary Data File 02. Data used to make Figure 2

Supplementary Data File 03. Data used to make Figure 3

Supplementary Data File 04. Data used to make Figure 4

## 12 Data Availability Statement

All data used in this manuscript can be found within the Supplemental Files provided.

## Notes

### Competing Interest Statement

The authors have declared no competing interest.

### Summary of Updates

Added ORCiD for one author

## References

Allen, M. B. (1959). Studies with cyanidium caldarium, an anomalously pigmented chlorophyte. Archiv. Mikrobiol. 32, 270–277. doi: 10.1007/BF00409348.

Almazroui, M., Nazrul Islam, M., Athar, H., Jones, P. D., and Rahman, M. A. (2012). Recent climate change in the Arabian Peninsula: annual rainfall and temperature analysis of Saudi Arabia for 1978-2009. Int. J. Climatol. 32, 953–966. doi: 10.1002/joc.3446.

AlSarmi, S. H., and Washington, R. (2014). Changes in climate extremes in the Arabian Peninsula: analysis of daily data: CHANGES EXTREMES OVER ARABIA. Int. J. Climatol. 34, 1329–1345. doi: 10.1002/joc.3772.

Cai, J., Lovatelli, A., Garrido Gamarro, E., Geehan, J., Lucente, D., Mair, G., et al. (2021). Seaweeds and microalgae: an overview for unlocking their potential in global aquaculture development. FAO doi: 10.4060/cb5670en.

Chen, Y., and Vaidyanathan, S. (2012). A simple, reproducible and sensitive spectrophotometric method to estimate microalgal lipids. Analytica Chimica Acta 724, 67–72. doi: 10.1016/j.aca.2012.02.049.

Chen, Y., and Vaidyanathan, S. (2013). Simultaneous assay of pigments, carbohydrates, proteins and lipids in microalgae. Analytica Chimica Acta 776, 31–40. doi: 10.1016/j.aca.2013.03.005.

Coward, T., Fuentes-Grünewald, C., Silkina, A., Oatley-Radcliffe, D. L., Llewellyn, G., and Lovitt, R. W. (2016). Utilising light-emitting diodes of specific narrow wavelengths for the optimization and co-production of multiple high-value compounds in Porphyridium purpureum. Bioresource Technology 221, 607–615. doi: 10.1016/j.biortech.2016.09.093.

de Freitas, B. B., Overmans, S., Medina, J. S., Hong, P.-Y., and Lauersen, K. J. (2023). Biomass generation and heterologous isoprenoid milking from engineered microalgae grown in anaerobic membrane bioreactor effluent. Water Research 229, 119486. doi: 10.1016/j.watres.2022.119486.

Ekendahl, S., Bark, M., Engelbrektsson, J., Karlsson, C.-A., Niyitegeka, D., and Strömberg, N. (2018). Energy-efficient outdoor cultivation of oleaginous microalgae at northern latitudes using waste heat and flue gas from a pulp and paper mill. Algal Research 31, 138–146. doi: 10.1016/j.algal.2017.11.007.

Ferdouse, F., Holdt, S. L., Smith, R., Murúa, P., and Yang, Z. (2018). The global status of seaweed production, trade and utilization. Available at: http://www.fao.org/in-action/globefish/publications/details-publication/en/c/1154074/.

Fick, S. E., and Hijmans, R. J. (2017). WorldClim 2: new 1-km spatial resolution climate surfaces for global land areas. Int. J. Climatol 37, 4302–4315. doi: 10.1002/joc.5086.

Fuentes-Grünewald, C., Ignacio Gayo-Peláez, J., Ndovela, V., Wood, E., Vijay Kapoore, R., and Anne Llewellyn, C. (2021). Towards a circular economy: A novel microalgal two-step growth approach to treat excess nutrients from digestate and to produce biomass for animal feed. Bioresource Technology 320, 124349. doi: 10.1016/j.biortech.2020.124349.

Greene, C. H., Scott-Buechler, C. M., Hausner, A. L. P., Johnson, Z. I., Lei, X. G., and Huntley, M. E. (2022). Transforming the future of marine aquaculture a circular economy approach. Oceanography 35. doi: 10.5670/oceanog.2022.213.

Guillard, R., R.,. L. ed. (1975). “Culture of phytoplankton for feeding marine invertebrates,” in Culture of phytoplankton for feeding marine invertebrates (Boston, MA: Springer US). doi: 10.1007/978-1-4615-8714-9.

Henkanatte-Gedera, S. M., Selvaratnam, T., Caskan, N., Nirmalakhandan, N., Van Voorhies, W., and Lammers, P. J. (2015). Algal-based, single-step treatment of urban wastewaters. Bioresource Technology 189, 273–278. doi: 10.1016/j.biortech.2015.03.120.

Kapoore, R. V., Huete-Ortega, M., Day, J. G., Okurowska, K., Slocombe, S. P., Stanley, M. S., et al. (2019). Effects of cryopreservation on viability and functional stability of an industrially relevant alga. Sci Rep 9, 2093. doi: 10.1038/s41598-019-38588-6.

Lefers, R. M., Tester, M., and Lauersen, K. J. (2020). Emerging Technologies to Enable Sustainable Controlled Environment Agriculture in the Extreme Environments of Middle East-North Africa Coastal Regions. Frontiers in Plant Science 11, 1–7. doi: 10.3389/fpls.2020.00801.

Matsuzaki, M., Misumi, O., Shin-i, T., Maruyama, S., Takahara, M., Miyagishima, S., et al. (2004). Genome sequence of the ultrasmall unicellular red alga Cyanidioschyzon merolae 10D. Nature 428, 653–657. doi: 10.1038/nature02398.

Mayhead, E., Silkina, A., Llewellyn, C., and Fuentes-Grünewald, C. (2018). Comparing Nutrient Removal from Membrane Filtered and Unfiltered Domestic Wastewater Using Chlorella vulgaris. Biology 7, 12. doi: 10.3390/biology7010012.

Minoda, A., Sakagami, R., Yagisawa, F., Kuroiwa, T., and Tanaka, K. (2004). Improvement of culture conditions and evidence for nuclear transformation by homologous recombination in a red alga, Cyanidioschyzon merolae 10D. Plant and Cell Physiology 45, 667–671. doi: 10.1093/pcp/pch087.

Miyagishima, S. Y., and Tanaka, K. (2021). The Unicellular Red Alga Cyanidioschyzon merolae - The Simplest Model of a Photosynthetic Eukaryote. Plant and Cell Physiology 62, 926–941. doi: 10.1093/pcp/pcab052.

Nirmalakhandan, N., Selvaratnam, T., Henkanatte-Gedera, S. M., Tchinda, D., Abeysiriwardana-Arachchige, I. S. A., Delanka-Pedige, H. M. K., et al. (2019). Algal wastewater treatment: Photoautotrophic vs. mixotrophic processes. Algal Research 41, 101569. doi: 10.1016/j.algal.2019.101569.

Rahman, D. Y., Sarian, F. D., van Wijk, A., Martinez-Garcia, M., and van der Maarel, M. J. E. C. (2017). Thermostable phycocyanin from the red microalga Cyanidioschyzon merolae, a new natural blue food colorant. J Appl Phycol 29, 1233–1239. doi: 10.1007/s10811-016-1007-0.

Ras, M., Steyer, J.-P., and Bernard, O. (2013). Temperature effect on microalgae: a crucial factor for outdoor production. Rev Environ Sci Biotechnol 12, 153–164. doi: 10.1007/s11157-013-9310-6.

Rumin, J., Nicolau, E., Gonçalves de Oliveira Junior, R., Fuentes-Grünewald, C., Flynn, K. J., and Picot, L. (2020). A Bibliometric Analysis of Microalgae Research in the World, Europe, and the European Atlantic Area. Marine Drugs 18, 79. doi: 10.3390/md18020079.

Selvaratnam, T., Henkanatte-Gedera, S. M., Muppaneni, T., Nirmalakhandan, N., Deng, S., and Lammers, P. J. (2016). Maximizing recovery of energy and nutrients from urban wastewaters. Energy 104, 16–23. doi: 10.1016/j.energy.2016.03.102.

Silkina, A., Ginnever, N. E., Fernandes, F., and Fuentes-Grünewald, C. (2019). Large-Scale Waste Bio-Remediation Using Microalgae Cultivation as a Platform. Energies 12, 2772. doi: 10.3390/en12142772.

Singh, M., and Das, K. C. (2014). “Low cost nutrients for algae cultivation,” in Algal biorefineries: Volume 1: Cultivation of cells and products, eds. R. Bajpai, A. Prokop, and M. Zappi (Dordrecht: Springer Netherlands), 69–82. doi: 10.1007/978-94-007-7494-0_3.

Toda, K., Takahashi, H., Itoh, R., and Kuroiwa, T. (1995). DNA Contents of Cell Nuclei in Two Cyanidiophyceae: Cyanidioschyzon merolae and Cyanidium caldarium Forma A. cytologia 60, 183–188. doi: 10.1508/cytologia.60.183.11

